# Multivariate links between the developmental timing of adversity exposure and white matter tract integrity in adulthood

**DOI:** 10.1101/2023.11.12.566271

**Authors:** Lucinda M. Sisk, Taylor J. Keding, Emily M. Cohodes, Sarah McCauley, Jasmyne C. Pierre, Paola Odriozola, Sahana Kribakaran, Jason T. Haberman, Sadie J. Zacharek, H. R. Hodges, Camila Caballero, Gillian Gold, Audrey Y. Huang, Ashley Talton, Dylan G. Gee

**Affiliations:** Department of Psychology, Yale University, New Haven, CT, USA; Department of Psychiatry, University of Wisconsin School of Medicine & Public Health, Madison, WI, USA; Department of Psychology, The City College of New York, New York, NY, USA; Department of Psychology, University of California Los Angeles, Los Angeles, CA, USA; Interdepartmental Neuroscience Program, Yale School of Medicine, New Haven, CT, USA; Department of Brain and Cognitive Sciences, Massachusetts Institute of Technology, Cambridge, MA, USA; Institute of Child Development, University of Minnesota, Minneapolis, MN, USA

## Abstract

Early-life adversity is pervasive worldwide and represents a potent risk factor for increased mental health burden across the lifespan. However, there is substantial individual heterogeneity in associations between adversity exposure, neurobiological changes, and mental health problems. Accounting for key features of adversity such as the developmental timing of exposure may clarify associations between adversity, neurodevelopment, and mental health. The present study leverages sparse canonical correlation analysis to characterize modes of covariation between age of adversity exposure and the integrity of white matter tracts throughout the brain in a sample of 107 adults. We find that adversity exposure during middle childhood (ages 5-6 and 8-9 in particular) is consistently linked with alterations in white matter tract integrity, such that tracts supporting sensorimotor functions display higher integrity in relation to adversity exposure while tracts supporting cortico-cortical communication display lower integrity. Further, latent patterns of tract integrity linked with adversity experienced across preschool age and middle childhood (ages 4-9) were associated with trauma-related symptoms in adulthood. Our findings underscore that adversity exposure may differentially affect white matter in a function- and developmental-timing specific manner and suggest that adversity experienced between ages 4-9 may shape the development of global white matter tracts in ways that are relevant for adult mental health.

## Introduction

Up to 60% of youth worldwide are exposed to adverse childhood experiences^1^. Childhood adversity is a potent predictor of worsened mental health burden later in life^2^, yet, at the individual level, there is substantial variability in the onset and severity of mental health disorders following adversity exposure. Influential theoretical models posit that specific aspects of adversity exposure, such as the type, timing, severity, and individuals’ perceptions of the events, may contribute to this heterogeneity in outcomes^3–6^. Given ongoing maturation of neural systems supporting cognition, emotion, and sensory processing and integration throughout childhood and adolescence, the developmental timing of exposure is posited to uniquely modulate the impacts of adversity on later mental health^7,8^. Identifying specific developmental stages at which adversity exposure is likely to confer elevated risk for mental health-related neurobiological changes represents a crucial step toward mechanistically defining relations between adversity exposure and mental health burden.

Clarifying when and how environmental experiences such as adversity shape neural structure, and whether such alterations are linked with later mental health problems, requires consideration of the state of the developing brain. The brain matures along a sensorimotor to association axis, such that networks supporting sensorimotor input, processing, and activity, mature earlier in childhood, while higher-order association and transmodal integration networks mature later in adolescence and early adulthood^9,10^. Given that brain structure and function develop in a modular, circuit-specific manner^11,12^, the precise state of the developing brain at the time of adversity exposure may partially determine how the exposure affects subsequent neurodevelopment and behavior^7,8,13–17^. Typical neurodevelopment includes windows of heightened experience-dependent plasticity, during which circuits display increased sensitivity to environmental input pertinent to their function^18,19^. Thus, circuits may be particularly vulnerable to function-relevant adverse events that intersect with these periods of heightened sensitivity, resulting in disproportionate changes when compared to the same event experienced during a different developmental period^14^. In this manner, circuit-specific, experience-dependent neuroplasticity may represent a mechanism whereby adverse experiences differentially impact individuals’ mental health depending on the developmental timing of exposure, contributing to cascading individual differences across development.

Empirical studies investigating associations between the developmental timing of adversity exposure and the brain have predominantly focused on neural regions known to be stress-sensitive, such as the hippocampus, amygdala, and prefrontal cortex^8,20^. Such work has provided cross-species evidence for distinct periods of increased sensitivity to adversity exposure at different stages. For example, amygdala structure and function demonstrate marked associations with adversity exposure during the juvenile period in rodents and pre-adolescence in youth^21,22^. White matter tracts supporting communication among stress-sensitive regions have also received significant attention. Indeed, exposure to adversity during the pre-adolescent period is associated with reduced volume of the corpus callosum, the primary white matter tract supporting interhemispheric cortical communication^23,24^, while severe neglect in early childhood is associated with brain-wide microstructural alterations^25^. More specifically, neglect in early childhood is associated with lower integrity of the corpus callosum, frontostriatal tracts, and limbic tracts, but higher integrity of tracts involved in sensorimotor processing^25^. Together, these findings suggest a negative association between adversity and white matter integrity in tracts supporting higher-order processing, such as the corpus callosum, superior longitudinal fasciculus, and uncinate fasciculus, and a positive association between adversity exposure and tracts supporting sensorimotor functions, particularly during early childhood and pre-adolescence. However, no studies to our knowledge have systematically examined the relative associations between adversity across development and white matter tract integrity across the brain.

Due to the critical role of white matter in conducting information across and between developing circuits, white matter tract integrity is of particularly interest when considering intersecting effects of adversity and developmental timing on brain circuitry. Building on foundational research identifying critical periods for visual perception in non-human animal^19^, current understanding of sensitive and critical periods of brain development includes multiple cellular mechanisms that facilitate increased neuroplasticity^26^. For example, preferential myelination of axonal pathways that exhibit greater recruitment is an experience-dependent process by which neuroplasticity enables adaptation to the environment^27^. Indeed, myelination is a critical mechanism that constrains neuroplasticity, particularly during adolescence^10,14,28^. Probing whether myelination contributes to adversity-associated alterations in white matter integrity––and whether this varies by the timing of adversity exposure––will facilitate greater understanding of adaptive neural responses in the face of challenging experiences^18,29^.

In the present study, we leverage a data-driven approach to examine covariation between adversity exposure at specific developmental stages and anatomical properties of white matter tracts in adulthood, as well as whether these patterns of covariation are associated with mental health symptoms. Using sparse canonical correlation analysis (sCCA)^30^, a multivariate method that permits simultaneous evaluation of two high-dimensional datasets, we examine associations between forty-three brain-wide white matter tracts and the developmental timing of adversity exposure. To identify patterns across metrics of tract integrity that may provide complementary insight into underlying mechanisms, we utilized three independent models to evaluate links between adversity and generalized fractional anisotropy (GFA)^31^, quantitative anisotropy (QA)^32^, and radial diffusivity (RD)^33^. Whereas GFA is thought to measure multiple axonal properties at the voxel level, such as fiber organization, crossing, and myelination, QA indexes these properties for the primary fiber orientation contained within the voxel^32^. For both measures, lower GFA and QA would be associated with reduced tract integrity. RD is sensitive to demyelination, such that higher RD is associated with lower myelin content^34^. Informed by a rich literature on the neurodevelopmental effects of early childhood adversity^25^, we hypothesized that greater exposure to adversity between the ages of 0-3 would be linked with widespread decrements in tract integrity as measured by GFA and QA, with the most pronounced negative associations observed in the uncinate fasciculus and corpus callosum. Second, in line with recent evidence that adolescence may represent a period of heightened plasticity for association cortex neurodevelopment^10,35^, we hypothesized that a measure indexing myelin content (RD) in tracts connecting regions of association cortex (e.g., the superior longitudinal fasciculus, the inferior longitudinal fasciculus) would be negatively linked with exposure to adversity during adolescence (ages 11-15). Finally, we hypothesized that variates primarily representing adversity in early childhood and integrity of the uncinate fasciculus and corpus callosum would both be associated with mental health symptoms in adulthood.

## Results

### sCCA Model Fitting

We utilized sCCA to identify links between the developmental timing of adversity exposure during childhood and white matter integrity in adulthood across forty-three tracts in the brain. Covariates were regressed from the adversity and white matter tract data separately prior to submission to the model (see Methods). Specifically, for each of the three white matter metrics of interest (GFA, QA, and RD), we fitted separate sCCA models across 10,000 iterations, with the number of modes (i.e., pairs of correlated variates) set to the number of adversity measures submitted to the model (19 modes). We aimed to identify (a) whether mode correlation strength differed significantly between the unshuffled (fitted) and shuffled (null) models, and (b) which modes explained the greatest proportion of total covariance. Across all models, the first mode explained the greatest amount of covariance (**Figure 1a**) and had the strongest correlation between mode variates (white matter and adversity variates, respectively). Thus, we opted to examine only the first mode for each of the GFA, QA, and RD models. For each of these models, permutation testing demonstrated that the first mode (**Figure 1b**) had a greater correlation strength than did the first mode of the null model (all *p*s < .05 after False Discovery Rate [FDR] correction)^36^.

**Figure 1.**
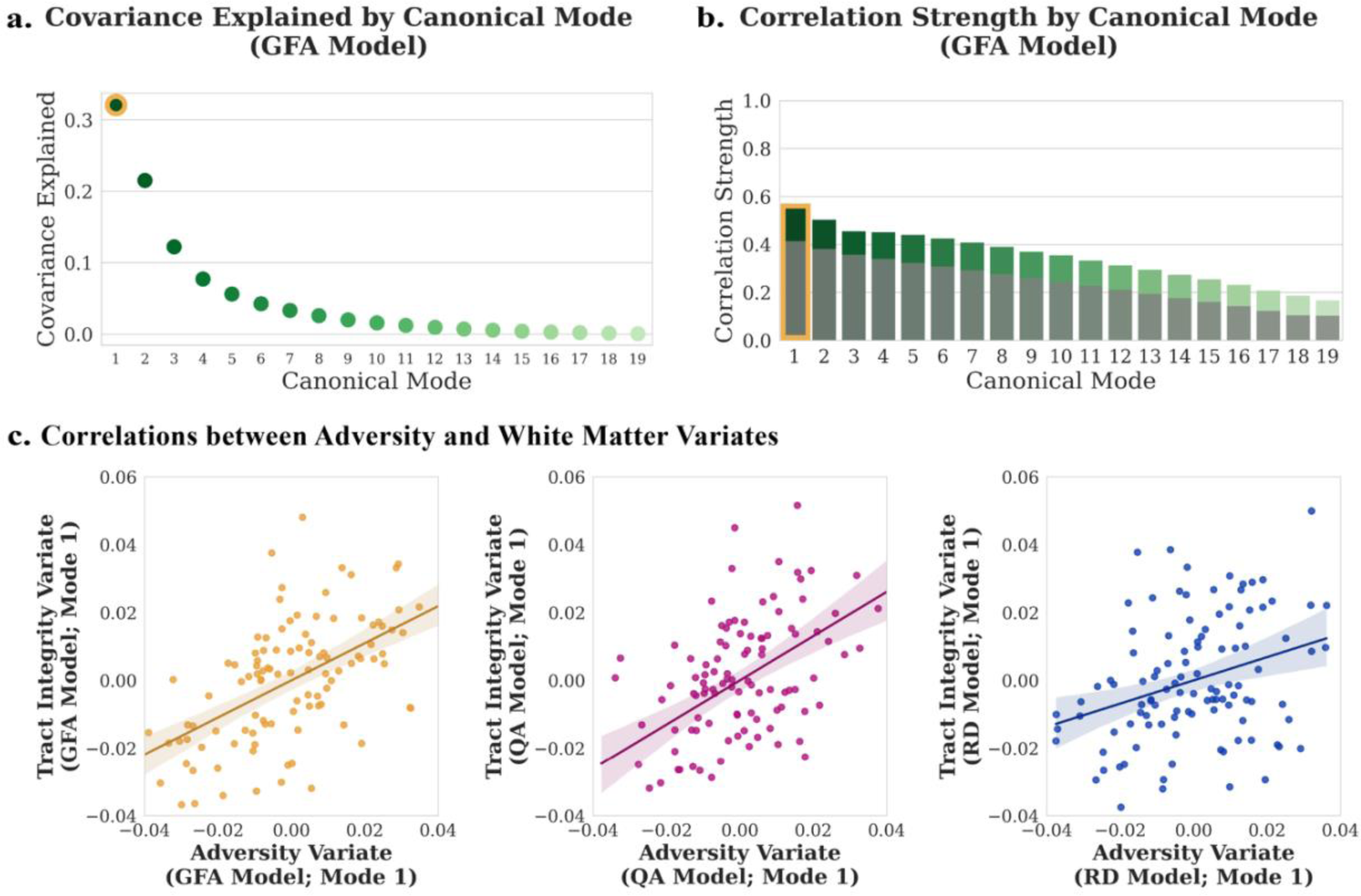
Model covariance explained and correlation strengths, exemplified in the GFA model. **a.** Median covariance explained by each canonical mode over 10,000 iterations of resampling. **b.** Median correlation strengths between the variates of the 19 canonical modes over 10,000 iterations of resampling. The median correlation strengths between variates of the shuffled (null) model over 10,000 iterations of resampling are overlaid in gray upon the median correlation strengths between variates of the unshuffled model (green). **c.** Associations between the adversity and white matter variates of each of the three models (GFA, QA, RD). GFA = generalized fractional anisotropy, QA = quantitative anisotropy, RD = radial diffusivity.

### GFA Model

Next, we tested whether variables loading onto model variates (i.e., adversity and white matter contributors to a given mode) in unshuffled models differed statistically in magnitude from the variables loading onto model variates in shuffled models using Wilcoxon signed-rank tests^37^. For the adversity variate of the first mode of the GFA model, all adversity exposure variables differed significantly between the unshuffled model loadings and the shuffled model loadings (Wilcoxon tests, all *p*s_FDR_ < .05; **Table 1**), with the exception of exposure to adversity at ages 10-11, 11-12, 12-13, 14-15, and 16-17. Similarly, all white matter tract loadings except for the right inferior longitudinal fasciculus differed significantly from the white matter tract loadings on the shuffled model (Wilcoxon tests, all *p*s_FDR_ < .05; **Table 2**). To identify which variables were most strongly represented in the first variate, we examined the median loading values for variables loading onto the adversity and white matter variates of the first mode across 10,000 bootstrapped model fits, excluding non-significant variables. Because variables could load both positively and negatively onto the variates, we used the absolute value of the median loading as an index of a given variable’s importance and examined the top 25% most important variables (**Figure 2a**). The top loadings for the adversity variate were adversity experienced during ages 8-9 and 5-6 years, followed by adversity experienced during ages 1-2, 0-1, and 7-8 years, respectively. All adversity exposure variables loaded negatively onto the adversity variate. In the white matter variate, loadings representing the integrity of the bilateral corticospinal tracts, fronto-pontine tracts, and parietal occipital-pontine tracts were strongest, followed by the right superior longitudinal fasciculus III tract, bilateral superior thalamic radiation tracts, and the corpus callosum isthmus and splenium. The superior longitudinal fasciculus and corpus callosum segments loaded positively onto the variate, whereas the corticospinal tracts, cortico-pontine tracts, and thalamo-cortical tracts loaded negatively onto the variate (**Figure 2a**).

**Figure 2.**
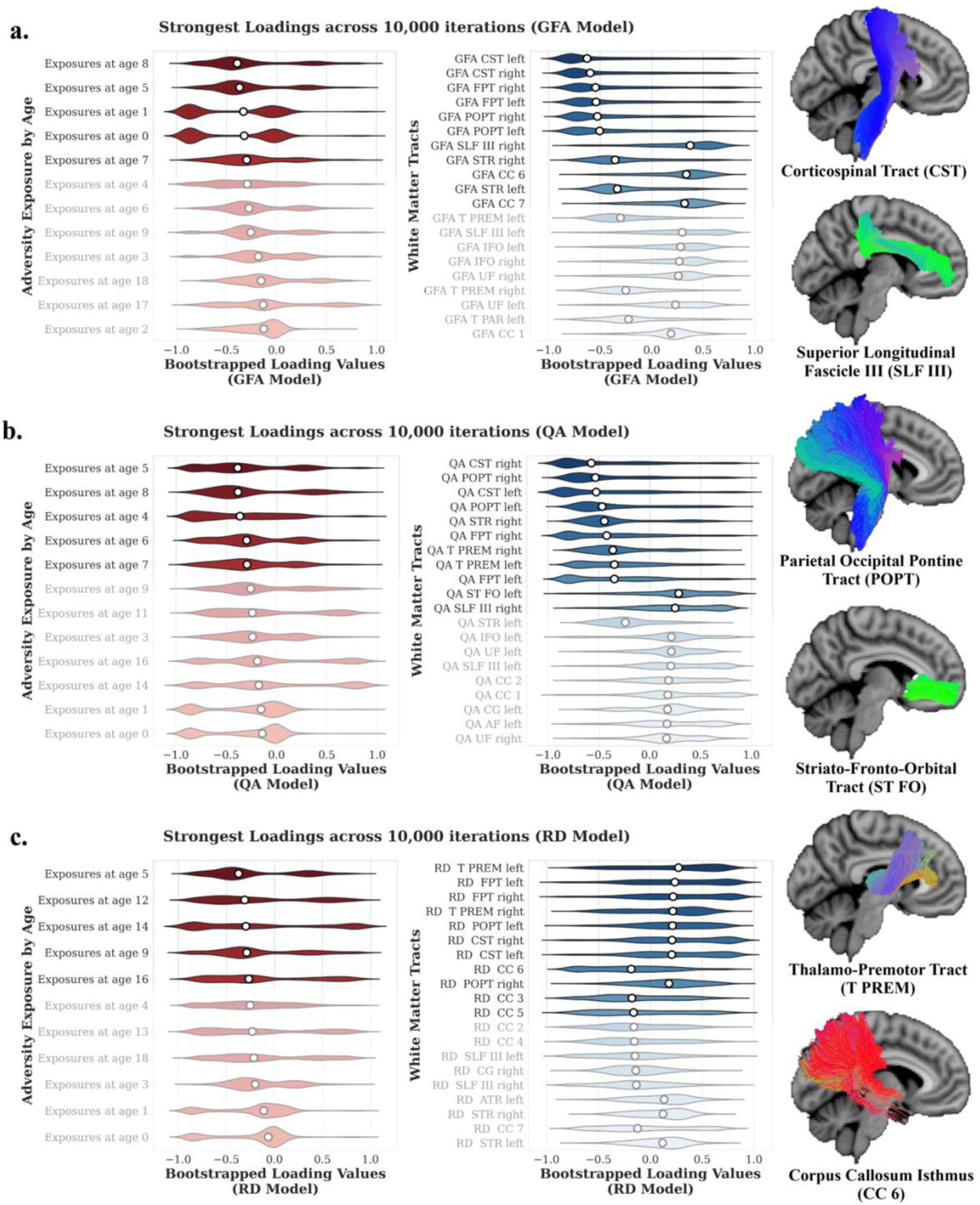
Top loadings for adversity (left) and tract integrity (right). The median loading value for each variable (white circle) is plotted on top of the distribution of bootstrapped loading values for that variable. The top 25% of tracts are displayed, with lower loadings greyed out. **a.** Loadings from the GFA model. **b.** Loadings from the QA model. **c.** Loadings from the RD model. Segmented tract images from a representative participant are plotted on the far right. GFA = generalized fractional anisotropy, QA = quantitative anisotropy. RD = radial diffusivity. AF = arcuate fasciculus, ATR = anterior thalamic radiation, CC 1 = corpus callosum rostrum, CC 2 = corpus callosum genu, CC 3 = corpus callosum rostral body (premotor), CC 4 = corpus callosum anterior midbody (primary motor), CC 6 = corpus callosum isthmus, CC 7 = corpus callosum splenium, CG = cingulum cingulate, CST = corticospinal tract, FPT = fronto-pontine tract, IFO = inferior fronto-occipital fasciculus, POPT = parietal occipital pontine tract, SLF III = superior longitudinal fasciculus III, ST FO = striato-fronto-orbital tract, STR = superior thalamic radiation, T PREM = thalamo-premotor tract, UF = uncinate fasciculus.

**Table 1.** Loading significance for variables on the adversity variate of mode 1 across models.

| Variable | GFA Model: Wilcoxon Stat | GFA Model: p-value (FDR-corrected) | QA Model: Wilcoxon Stat | QA Model: p-value (FDR-corrected) | RD Model: Wilcoxon Stat | RD Model: p-value (FDR-corrected) |
| --- | --- | --- | --- | --- | --- | --- |
| Exposure at age 0 | 12745370 | <.001 | 17568324 | <.001 | 22205637 | <.001 |
| Exposure at age 1 | 12522215 | <.001 | 17773747 | <.001 | 21913992 | <.001 |
| Exposure at age 2 | 17784643 | <.001 | 18408360 | <.001 | 24571131 | 0.197 |
| Exposure at age 3 | 22841243 | <.001 | 23643819 | <.001 | 23292725 | <.001 |
| Exposure at age 4 | 22077269 | <.001 | 19137768 | <.001 | 24206530 | 0.011 |
| Exposure at age 5 | 22660616 | <.001 | 20607266 | <.001 | 23803166 | <.001 |
| Exposure at age 6 | 23606428 | <.001 | 21904761 | <.001 | 24968918 | 0.907 |
| Exposure at age 7 | 22864874 | <.001 | 22573744 | <.001 | 24928093 | 0.89 |
| Exposure at age 8 | 23781817 | <.001 | 23921343 | <.001 | 24606163 | 0.215 |
| Exposure at age 9 | 24202114 | 0.009 | 24228374 | 0.009 | 24309201 | 0.028 |
| Exposure at age 10 | 24637731 | 0.231 | 24741335 | 0.366 | 24595949 | 0.215 |
| Exposure at age 11 | 24774864 | 0.454 | 23824382 | <.001 | 24721272 | 0.392 |
| Exposure at age 12 | 24448971 | 0.07 | 24683272 | 0.284 | 23753516 | <.001 |
| Exposure at age 13 | 24307603 | 0.022 | 24436367 | 0.056 | 23921387 | <.001 |
| Exposure at age 14 | 24487460 | 0.088 | 23419908 | <.001 | 23740089 | <.001 |
| Exposure at age 15 | 22080824 | <.001 | 22083423 | <.001 | 24956515 | 0.907 |
| Exposure at age 16 | 24942978 | 0.837 | 24042522 | 0.001 | 23966207 | 0.001 |
| Exposure at age 17 | 23726903 | <.001 | 22216378 | <.001 | 24528233 | 0.159 |
| Exposure at age 18 | 24249047 | 0.013 | 22460632 | <.001 | 23785363 | <.001 |

**Table 2.** Loading significance for variables on the white matter variate of mode 1 across models. AF = arcuate fasciculus, ATR = anterior thalamic radiation, CC 1 = corpus callosum rostrum, CC 2 = corpus callosum genu, CC 3 = corpus callosum rostral body (premotor), CC 4 = corpus callosum anterior midbody (primary motor), CC 6 = corpus callosum isthmus, CC 7 = corpus callosum splenium, CG = cingulum cingulate, CST = corticospinal tract, FPT = fronto-pontine tract, IFO = inferior fronto-occipital fasciculus, OR=optic radiation, POPT = parietal occipital pontine tract, SLF I = superior longitudinal fasciculus I, SLF II = superior longitudinal fasciculus II, SLF III = superior longitudinal fasciculus III, ST FO = striato-fronto-orbital tract, STR = superior thalamic radiation, T PAR = thalamo-parietal tract, T PREM = thalamo-premotor tract, UF = uncinate fasciculus.

| Variable | GFA Model: Wilcoxon Stat | GFA Model: p-value (FDR-corrected) | QA Model: Wilcoxon Stat | QA Model: p-value (FDR-corrected) | RD Model: Wilcoxon Stat | RD Model: p-value (FDR-corrected) |
| --- | --- | --- | --- | --- | --- | --- |
| AF left | 19755762 | <.001 | 15835923 | <.001 | 20431837 | <.001 |
| AF right | 16429076 | <.001 | 21442334 | <.001 | 24328362 | 0.021 |
| ATR left | 20302961 | <.001 | 24228312 | 0.008 | 17841522 | <.001 |
| ATR right | 17968813 | <.001 | 23611604 | <.001 | 20193716 | <.001 |
| CC 1 | 14008988 | <.001 | 14179620 | <.001 | 20629020 | <.001 |
| CC 2 | 18480349 | <.001 | 14769985 | <.001 | 18088455 | <.001 |
| CC 3 | 16949427 | <.001 | 20258529 | <.001 | 18596247 | <.001 |
| CC 4 | 20162473 | <.001 | 21025428 | <.001 | 19344047 | <.001 |
| CC 5 | 21002177 | <.001 | 24010030 | 0.001 | 20650510 | <.001 |
| CC 6 | 11685113 | <.001 | 18414716 | <.001 | 15846890 | <.001 |
| CC 7 | 10957243 | <.001 | 14876370 | <.001 | 22114656 | <.001 |
| CG left | 13400467 | <.001 | 17230012 | <.001 | 17978803 | <.001 |
| CG right | 13656752 | <.001 | 20601903 | <.001 | 15351082 | <.001 |
| CST left | 10252111 | <.001 | 11461585 | <.001 | 15175168 | <.001 |
| CST right | 9924477 | <.001 | 10597033 | <.001 | 14693124 | <.001 |
| FPT left | 11645178 | <.001 | 12876038 | <.001 | 15710949 | <.001 |
| FPT right | 11969612 | <.001 | 11999876 | <.001 | 15438342 | <.001 |
| IFO left | 13630342 | <.001 | 16693919 | <.001 | 24894190 | 0.708 |
| IFO right | 14882295 | <.001 | 20708964 | <.001 | 21442845 | <.001 |
| ILF left | 17270446 | <.001 | 18844120 | <.001 | 19394430 | <.001 |
| ILF right | 24796450 | 0.475 | 24091696 | 0.002 | 21589152 | <.001 |
| OR left | 18296967 | <.001 | 19059172 | <.001 | 21198967 | <.001 |
| OR right | 23668045 | <.001 | 24620905 | 0.186 | 24435385 | 0.051 |
| POPT left | 10099285 | <.001 | 12240574 | <.001 | 14399824 | <.001 |
| POPT right | 9419399 | <.001 | 11055311 | <.001 | 15234551 | <.001 |
| SLF I left | 19978984 | <.001 | 22682165 | <.001 | 22678797 | <.001 |
| SLF I right | 23845083 | <.001 | 19373901 | <.001 | 21489828 | <.001 |
| SLF II left | 15570684 | <.001 | 22791342 | <.001 | 24384809 | 0.034 |
| SLF II right | 19431927 | <.001 | 22023376 | <.001 | 21948828 | <.001 |
| SLF III left | 11465218 | <.001 | 13405628 | <.001 | 16542897 | <.001 |
| SLF III right | 11979376 | <.001 | 13570599 | <.001 | 16303655 | <.001 |
| ST FO left | 18406367 | <.001 | 14284547 | <.001 | 24315003 | 0.019 |
| ST FO right | 18077478 | <.001 | 15073037 | <.001 | 22020476 | <.001 |
| ST PREM left | 19812882 | <.001 | 18766639 | <.001 | 24157897 | 0.004 |
| ST PREM right | 19746843 | <.001 | 19036842 | <.001 | 24042437 | 0.001 |
| STR left | 12693033 | <.001 | 14379468 | <.001 | 18347995 | <.001 |
| STR right | 13194333 | <.001 | 11731548 | <.001 | 18847434 | <.001 |
| T PAR left | 15178042 | <.001 | 20497577 | <.001 | 19335684 | <.001 |
| T PAR right | 16682189 | <.001 | 15980436 | <.001 | 21649649 | <.001 |
| T PREM left | 12242526 | <.001 | 12137483 | <.001 | 16830558 | <.001 |
| T PREM right | 13119088 | <.001 | 11922357 | <.001 | 18391878 | <.001 |
| UF left | 13628185 | <.001 | 15255798 | <.001 | 23759828 | <.001 |
| UF right | 14395946 | <.001 | 16034894 | <.001 | 22750895 | <.001 |

### QA Model

For the QA model, all adversity exposure variables differed significantly between the unshuffled model loadings and the shuffled model loadings (Wilcoxon tests; all *p*s_FDR_ < .05; **Table 1**), with the exception of exposure at ages 10-11, 12-13, and 13-14 years. All white matter tract loadings except for the right optic radiation also differed significantly between the white matter tract loadings on the unshuffled model and the shuffled model loadings (Wilcoxon tests; all *p*s_FDR_ < .05; **Table 2**). We next examined the median loadings of the first variate of the QA model. The top loadings for the adversity variate were adversity experienced from ages 5-6 and 8-9 years, followed by adversity experienced during ages 4-5, 6-7, and 7-8 years, respectively. All adversity exposure variables loaded negatively onto the adversity variate. In the white matter variate, integrity of the bilateral corticospinal tracts, parietal occipital pontine tracts, right superior thalamic radiation, fronto-pontine tracts, and bilateral thalamo-premotor tracts loaded negatively onto the variate. The left striato-fronto-orbital tract and right superior longitudinal fasciculus III tract loaded positively onto the variate (**Figure 2b**).

### RD Model

For the RD model, all adversity exposure variables differed significantly between the unshuffled model loadings and the shuffled model loadings (Wilcoxon tests; all *ps*_FDR_ < .05; **Table 1**), with the exception of exposure at ages 2-3, 6-7, 7-8, 8-9, 10-11, 11-2, 15-16, and 17-18 years. All white matter tract loadings except for left inferior fronto-occipital fasciculus and the right optic radiation differed significantly from the white matter tract loadings on the shuffled model (Wilcoxon tests; all *p*s_FDR_ < .05, **Table 2**). We next examined the median loadings of the first variate of the RD model. The first adversity exposure loading for this model partially paralleled the GFA and QA models, with adversity exposure at ages 5-6 representing the top loading. However, rather than being followed by exposure in early and middle childhood, for the RD model, adversity exposure from ages 12-13, and 14-15, 9-10, and 16-17, respectively, represented the next strongest loadings. All adversity exposure variables loaded negatively onto the adversity variate. For the white matter variate, bilateral thalamo-premotor tracts, fronto-pontine tracts, corticospinal tracts, and parietal occipito-pontine tracts loaded positively, while the corpus callosum isthmus, genu, and rostral body loaded negatively (**Figure 2c**).

### Associations with Mental Health Symptoms

Next, we tested whether variates representing adversity exposure and white matter integrity, respectively, were associated with mental health symptoms. We assessed two measures of symptoms: total mental health symptoms derived from the Adult Self Report scale (ASR)^38^, and trauma-related symptoms derived from the Trauma Symptom Checklist (TSC-40)^39^. For each measure of symptoms, two separate Ordinary Least Squares (OLS) models tested (1) whether adversity variates from the GFA, QA, and RD sCCA models were associated with mental health symptoms, and (2) whether white matter variates from the GFA, QA, and RD sCCA models were associated with mental health symptoms. We used FDR correction to account for multiple comparisons (i.e., the two symptom measures)^36^. We found that the white matter variate from the QA model was associated with trauma-related symptoms above and beyond variates from the GFA and RD models (*t*(90) = −3.611, *β* = −0.546, *p_FDR_ =* .001); **Figure 3**). None of the variates representing adversity exposure by age were associated with either measure of symptoms (all *p*s > .05).

**Figure 3.**
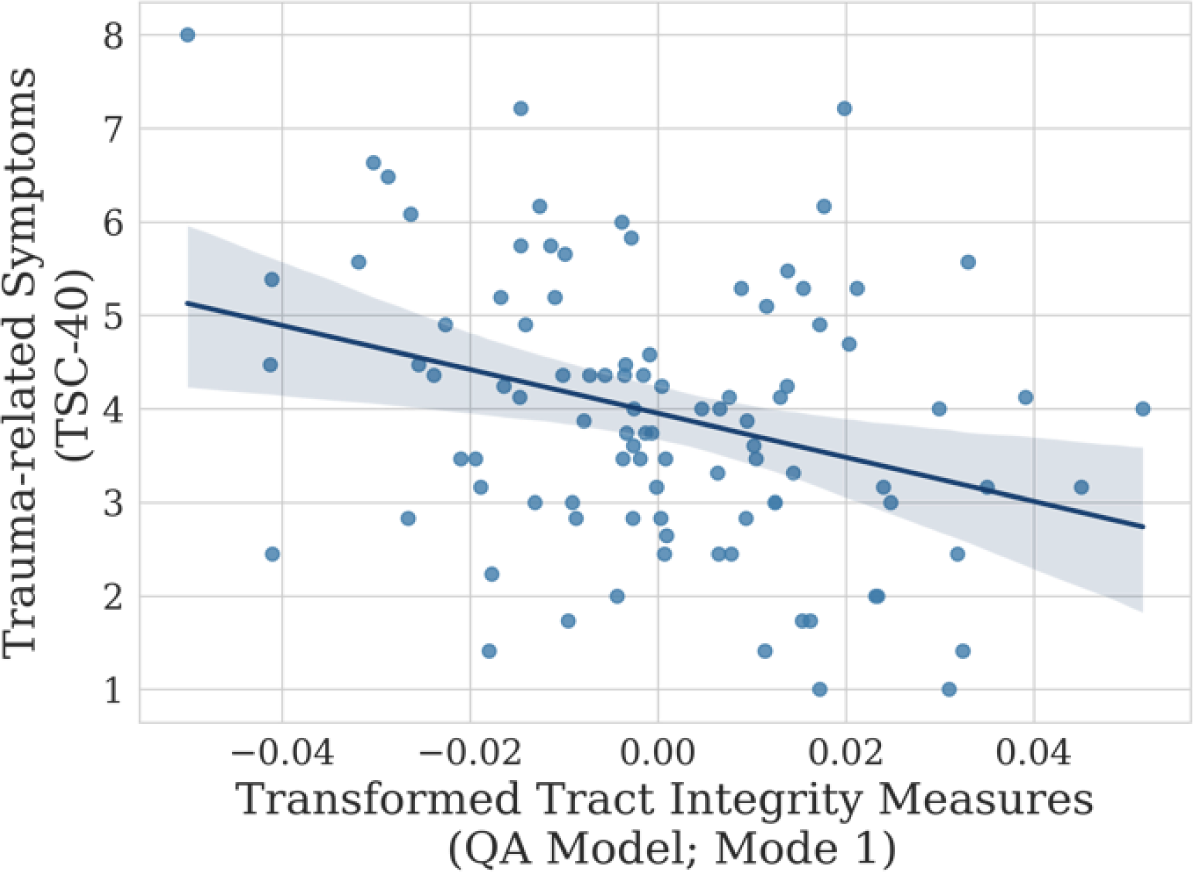
Associations between trauma-related symptoms and transformed values of the white matter variate from mode 1 of the QA model averaged across 10,000 iterations of resampling. QA = quantitative anisotropy.

### Sensitivity Analyses

We conducted several sensitivity analyses to probe the robustness of our findings when controlling for additional variables. First, given broad evidence that socioeconomic status is a proxy for experiences that may shape brain development^40,41^ and may covary with measures of adversity such that children who grow up in households with lower socioeconomic status are at heightened risk of experiencing adversity^13,42^, we tested whether our models might be identifying variates with variability related to socioeconomic status. For each of the three models (GFA, QA, and RD) we tested separately whether the variates of the first mode were correlated with combined family income or years of education. None of the variates displayed a meaningful correlation with income nor education (all *p*s > .05). Next, we tested whether including combined family income and years of education as covariates in our OLS models examining associations between model variates and trauma-related symptoms altered the findings. Including income and education as covariates in the white matter tract integrity model did not affect the significance of the association between trauma-related symptoms and the white matter variate of the first mode of the QA model (*t*(90) = −3.450, *β* = −0.521, p = .001). We also conducted follow-up OLS models to test whether any of the individual white matter tract variables in the top 25% of the strongest loadings were independently associated with trauma-related symptoms. No tracts were significantly associated with trauma-related symptoms (all *p*s > .05).

## Discussion

Adversity experienced during development can have profound and lasting effects on the brain and mental health; however, there is substantial heterogeneity in outcomes following adversity. In the present study, we leveraged a data-driven approach to investigate modes of covariation between the developmental timing of adversity and white matter tract integrity in adulthood. We observed a systematic pattern of covariance between white matter tracts and adversity, such that greater adversity exposure in middle childhood was linked with higher integrity of white matter tracts subserving sensorimotor functions (such as the corticospinal tract and corticopontine tracts) but lower integrity of white matter tracts subserving cortico-cortical communication (such as the corpus callosum and superior longitudinal fasciculus). In both GFA and QA models, adversity exposure between the ages of 5-6 and 8-9 was linked with lower integrity of cortico-cortical tracts and higher integrity of sensorimotor tracts. The RD model demonstrated the opposite pattern, such that results were consistent with the differential properties that RD indexes. That is, exposure between the ages of 5-6 was linked with higher RD in cortico-cortical tracts, suggesting reduced myelin levels, and lower RD in sensorimotor tracts, suggesting higher myelin levels. As lower myelin content may underlie reductions in integrity measured by GFA and QA, together these three metrics demonstrate a consistent pattern. Additionally, the white matter variate derived from QA measures was negatively associated with trauma-related symptoms. Notably, the correlated adversity variate from the QA model was not associated with trauma-related symptoms, suggesting that whole-brain patterns of tract integrity may modulate links between adversity exposure and mental health. In a sensitivity analysis, we demonstrated that the tract variables comprising the loadings of this variate were not associated with trauma-related symptoms, suggesting that global patterns of tract integrity may be more relevant to mental health than localized alterations in specific tracts.

The finding that sensorimotor and cortico-cortical tracts show diverging relationships with exposure to adversity in middle childhood advances the field toward a clearer understanding of the whole-brain impacts of adversity and later mental health. While contrary to our first hypothesis that we would observe global decrements in white matter tract integrity in conjunction with early childhood adversity, these results suggest that adversity-associated, experience-dependent remodeling of structural connectivity may dovetail with patterns of neurodevelopment unfolding along the sensorimotor-association axis^10^. Evidence from the RD model, suggesting higher myelin content along sensorimotor tracts and lower myelin content along cortico-cortical tracts in association with adversity exposure in middle childhood and adolescence, implicates myelination as a key mechanism underlying these changes, partially supporting our second hypothesis. Indeed, divergent patterns of white matter integrity between tracts subserving sensorimotor functions versus cortico-cortical communication have been previously reported in association with adversity^25,43^, and the present results may provide a more nuanced, timing-related characterization of this pattern. Further, heightened sensory responsivity has been reported following early adversity exposure, and is related to mental health symptoms^44^. Our findings raise the possibility that such effects are underpinned by neurostructural changes, and that changes to sensory processing tracts may be understudied neuroanatomical correlates of adversity exposure. One possibility is that exposure to adversity, particularly in middle childhood, may activate experience-dependent myelination processes that prioritize enhanced structural connectivity supporting functions more critical to survival, such as motor activity and sensory perception and relay, and deprioritize structural connectivity supporting higher-order functions, such as complex cognitive integration and processing, among cortical association regions.

Despite the fact that the three metrics of tract integrity employed in this study (GFA, QA, and RD) are known to measure overlapping cellular properties^31–33^, they are computationally distinct and differentially characterize tract integrity. Thus, adversity exposure between the ages of 5-6 emerging across all three models as a top loading suggests that exposure during middle childhood may be associated with multiple mechanisms that govern white matter integrity. While we broadly interpret the findings across all models (i.e., links between adversity and white matter tract integrity) as being most strongly related to adversity in middle childhood due to ages 5-6 representing a strong loading across all three models and ages 8-9 representing a strong loading across GFA and QA models, it is worth noting that the models differ in their later loadings, potentially reflecting differential sensitivities of GFA, QA, and RD to underlying cellular features.

For instance, adversity exposure in adolescence comprised the 2^nd^-5^th^ loadings in the RD model, implicating myelination as a primary mechanism underlying neuroplasticity in adolescence. This finding aligns with recent work highlighting the role of myelin in constraining functional signal amplitude in adolescence^10^; however, further developmental work is needed to disentangle the precise timing of experience-dependent alterations to myelination trajectories across adolescence. In the GFA model, adversity exposure in early childhood (from ages 0-2) comprised the 3^rd^ and 4^th^ loadings. Possibly because of relatively low endorsement of adversity at these ages in the sample, these variables displayed a bimodal distribution across resampling iterations (**Figure 2b**), constraining our ability to draw inferences about effects associated with this particular developmental stage. Future studies utilizing larger samples and prospective measures of adversity exposure starting in infancy will be critical to identifying the relative impacts of adversity experienced during early versus middle childhood to patterns of white matter integrity, as will methodological advances that permit zero-inflated data (such as adversity exposure in early childhood) to be modeled more accurately.

Finally, the QA model displayed loadings representing adversity exposure experienced during preschool age and middle childhood (ages 4-9) across the top five loadings. Because the diffusion variate of this model was associated with trauma-related symptoms, these findings suggests that middle childhood (in contrast to our initial hypothesis of early childhood) may represent both a sensitive window during which adversity can “get under the skin” to shape white matter integrity, and a time when adversity exposure may increase the relative likelihood of experiencing later mental health problems. Together, these results also suggest that the increased specificity of QA in indexing directional fiber populations may yield metrics that better reflect circuit-specific changes, which in turn relate to mental health. Future work testing the hypothesis that middle childhood represents a sensitive window during which experiences of adversity confer greater risk for future mental health problems could inform when to intervene to optimally support youth mental health.

While the present study contributes to the field’s understanding of how the timing of adversity exposure across development may differentially shape white matter tracts and mental health symptoms, there are nevertheless important limitations that should be addressed in future research. First, the present study was conducted using a modestly-sized laboratory sample of participants who completed a series of time-intensive clinical interviews, resulting in rich information on adversity exposure during development^45^. This time-intensive phenotyping is critical to advancing the science of adversity exposure and brain development, yet practically constrains that the sample size we could feasibly collect. While we aimed to characterize our sample as robustly as possible by applying resampling and permutation testing^46^ and utilizing a multivariate approach^47^, it will be important to replicate these findings in a larger, more demographically diverse sample in order to evaluate robustness and generalizability. Further, despite having rich information on adversity exposure, sample size precluded us from further parsing these events by additional relevant dimensions of adversity, such as perceived severity, threat, deprivation, controllability, and predictability. Such features of adversity likely relate to subsequent neural and behavioral phenotypes^3–6^, and future work building upon the present findings by examining the intersection of these dimensions with developmental timing^48^ could meaningfully elucidate how specific dimensions of adversity may differentially relate to neurodevelopment, risk, and resilience at distinct ages. Finally, these findings should be interpreted in the context of reliance on retrospective report of adversity exposure. Participants may not have been able to accurately recall nor report all adverse events at each age, especially those experienced in early childhood^49^. While this limitation of retrospective report has been consistently noted^50,51^, evidence suggests that retrospective report nonetheless demonstrates stronger associations with mental health symptoms than more “objective” measures of adversity^52,53^. Future studies employing prospective, large-scale longitudinal studies that systematically query children’s adversity exposure at each age are needed to advance understanding of how adversity confers risk for mental health disorders, as well as elucidate how adversity at distinct stages of development shapes neurodevelopmental trajectories.

The present study identified multivariate links between the developmental timing of adversity, divergent patterning of white matter tract integrity, and trauma-related mental health symptoms. We characterized latent variates representing exposure to adversity in middle childhood (primarily during ages 5-6 and 8-9) and higher white matter integrity of sensorimotor tracts (such as the corticospinal tracts, parietal-occipital pontine tracts, fronto-pontine tracts, and thalamo-premotor tracts) and lower white matter integrity of cortico-cortical tracts (such as the superior longitudinal fasciculi, corpus callosum, striato-fronto-orbital tracts, and inferior fronto-occipital fasciculi). Convergent results across three diffusion metrics, including myelin-sensitive RD, suggest that experience-dependent changes to myelination patterns may underlie adversity-related alterations in white matter tracts. The latent variate representing white matter tract integrity measured by QA was associated with trauma-related symptoms, suggesting that the neural bases of clinical symptomatology may be distributed across the brain, rather than localized to a given tract or region. Taken together, these findings begin to elucidate the complex ways that adversity exposure may intersect with neurodevelopmental timing to differentially shape global white matter tracts across childhood and adolescence. Leveraging multivariate models to identify latent links between environmental exposures and brain development that have clinical relevance could ultimately inform future translational research aimed at refining interventions for youth exposed to adversity.

## Methods

### Participant Recruitment

*Participants.* Participants were 107 adults between the ages of 18 and 30 recruited from the broader New Haven, CT community. Participants included in the current study were right-handed and free of contraindications for an MRI scan (e.g., no braces or metal implants). Additional exclusion criteria are detailed in the **Supplement**. Study procedures were approved by the institutional review board at Yale University, and each participant provided informed consent in writing.

### Demographics

Participants reported their age at session, sex assigned at birth, race and ethnicity, years of education, and annual combined household income. Further demographic information on this sample can be found in **Table 3**.

**Table 3.** Demographic information for the sample (N=107) is shown for age at session, sex assigned at birth, gender identity, race and ethnicity, level of education, combined family income, total problems T-scores endorsed on the Adult Self Report (ASR) scale, trauma-related symptoms (TSC-40), diagnostic group, and site of scan acquisition. Chi-squared tests were used to evaluate differences between groups with and without TSC-40 data for ordinal and categorical data, and T-tests were used to evaluate differences for continuous data. Table created using the ‘arsenal’ package in R.

|  | **Not included in sensitivity analyses (N=17)** | **Included in sensitivity analyses (N=90)** | **Total (N=107)** | ***p*-value** |
| --- | --- | --- | --- | --- |
| **Age** |  |  |  | 0.458 |
| Mean (SD) | 23.385 (3.054) | 22.753 (3.232) | 22.854 (3.199) |  |
| Range | 18.048 - 29.556 | 18.125 - 30.829 | 18.048 - 30.829 |  |
| **Sex at Birth** |  |  |  | 0.296 |
| Female | 13 (76.5%) | 57 (63.3%) | 70 (65.4%) |  |
| Male | 4 (23.5%) | 33 (36.7%) | 37 (34.6%) |  |
| **Gender Identity** |  |  |  | 0.330 |
| Female | 13 (81.2%) | 56 (62.2%) | 69 (65.1%) |  |
| Male | 3 (18.8%) | 33 (36.7%) | 36 (34.0%) |  |
| Transgender | 0 (0.0%) | 1 (1.1%) | 1 (0.9%) |  |
| Missing Data | 1 | 0 | 1 |  |
| **Education** |  |  |  | 0.629 |
| High School | 1 (5.9%) | 13 (14.4%) | 14 (13.1%) |  |
| College | 12 (70.6%) | 57 (63.3%) | 69 (64.5%) |  |
| Graduate School | 4 (23.5%) | 20 (22.2%) | 24 (22.4%) |  |
| **Combined Income** |  |  |  | 0.784 |
| <\$5,000 | 1 (10.0%) | 4 (4.4%) | 5 (5.0%) | |
| \$5,000-\$11,999 | 1 (10.0%) | 3 (3.3%) | 4 (4.0%) | |
| \$12,000-\$14,999 | 1 (10.0%) | 4 (4.4%) | 5 (5.0%) | |
| \$15,000-\$24,999 | 1 (10.0%) | 8 (8.9%) | 9 (9.0%) | |
| \$25,000-\$34,999 | 0 (0.0%) | 7 (7.8%) | 7 (7.0%) | |
| \$35,000-\$49,999 | 0 (0.0%) | 8 (8.9%) | 8 (8.0%) | |
| \$50,000-\$74,999 | 3 (30.0%) | 21 (23.3%) | 24 (24.0%) | |
| \$75,000-\$99,999 | 1 (10.0%) | 5 (5.6%) | 6 (6.0%) | |
| \$100,000+ | 2 (20.0%) | 30 (33.3%) | 32 (32.0%) | |
| Missing Data | 7 | 0 | 7 |  |
| **Race and Ethnicity** |  |  |  | 0.201 |
| Asian | 5 (29.4%) | 16 (17.8%) | 21 (19.6%) |  |
| Black or African American | 3 (17.6%) | 10 (11.1%) | 13 (12.1%) |  |
| Hispanic or Latinx | 2 (11.8%) | 9 (10.0%) | 11 (10.3%) |  |
| Multiracial | 3 (17.6%) | 5 (5.6%) | 8 (7.5%) |  |
| Non-Hispanic White | 4 (23.5%) | 49 (54.4%) | 53 (49.5%) |  |
| Prefer not to answer | 0 (0.0%) | 1 (1.1%) | 1 (0.9%) |  |
| **ASR Total Problems T-Score** |  |  |  | **0.022** |
| Mean (SD) | 39.000 (9.381) | 44.933 (9.070) | 44.086 (9.306) |  |
| Range | 27.000 - 64.000 | 28.000 - 71.000 | 27.000 - 71.000 |  |
| Missing Data | 2 | 0 | 2 |  |
| **TSC-40 Symptoms** |  |  |  | 0.542 |
| Mean (SD) | 13.857 (9.063) | 16.833 (12.578) | 16.619 (12.345) |  |
| Range | 0.000 - 27.000 | 0.000 - 63.000 | 0.000 - 63.000 |  |
| Missing Data | 10 | 0 | 10 |  |
| **Diagnostic Group** |  |  |  | 0.137 |
| No diagnosis | 14 (82.4%) | 55 (61.1%) | 69 (64.5%) |  |
| Anxiety Diagnosis | 3 (17.6%) | 20 (22.2%) | 23 (21.5%) |  |
| Other Diagnosis | 0 (0.0%) | 15 (16.7%) | 15 (14.0%) |  |
| **Site** |  |  |  | 0.148 |
| Brain Imaging Center | 4 (23.5%) | 38 (42.2%) | 42 (39.3%) |  |
| Magnetic Resonance Research Center | 13 (76.5%) | 52 (57.8%) | 65 (60.7%) |  |

#### Interview and assessment of adversity exposure

Participants completed the Dimensional Inventory of Stress and Trauma Across the Lifespan (DISTAL) interview^45^. The DISTAL is broadly based on the structure of the University of California, Los Angeles, Post-Traumatic Stress Disorder Reaction Index interview^54^ and is designed to assess key dimensions of adverse exposure including age of exposure^45^. The DISTAL contains two subsections: a broad set of screening questions for potential exposure to multiple types of adversity at three levels of exposure (directly experiencing, witnessing, and learning about the event happening to a close person) and event-specific modules that query additional details about each event endorsed. Participants reported on their exposure to 24 distinct types of adverse events: serious accidental injury, illness/medical trauma, community violence, domestic violence, school violence or school emergency, physical assault, disaster, sexual abuse, physical abuse, neglect, psychological maltreatment and emotional abuse, impaired caregiving, sexual assault, kidnapping and abduction, terrorism, bereavement and witnessing death, caregiver separation, war and political violence, forced displacement, trafficking and sexual exploitation, bullying, attempted suicide, witnessing suicide, and verbal conflict. Screening items for exposure to each type of adversity listed above were written to capture exposure to a particular type of adversity at the three levels of exposure (i.e., directly experiencing, witnessing, and learning about the event happening to a close person; at least three screener questions were included for each type of event^45^.

For each type of event that was endorsed in the screening portion of the interview, participants were then asked to report on the cumulative list of ages at which they experienced a particular type of adversity (0-30, or 0 through the current age of participant, if current age less than 30). If multiple events of the same type were reported, age of exposure at each event was recorded. Interview data underwent extensive cleaning and entry by a team of trained research assistants, overseen by a clinical doctoral student and the principal investigator^45^. These cleaned data were then summed to derive a total count of adverse events at each year of age from ages 0-30 (or 0 through the current age of each participant). Reported events that did not include an age were excluded. See **Table S1** for descriptive statistics of these endorsements.

#### Mental health symptoms

Measures of total mental health symptoms were derived from the Adult Self Report scale (ASR)^38^. Participants completed the ASR prior to undergoing their scanning session, and their responses were coded using Achenbach System of Empirically Based Assessment software. Measures of total trauma-related symptomatology were derived from the Trauma Symptoms Checklist (TSC-40)^39^, which participants also completed prior to their scanning session.

#### Scanning session

Participants were scanned on a 3T Siemens Magnetom Prisma scanner (Siemens, Erlangen, Germany) using a 32-channel head coil. N=65 participants were scanned at the Yale University Magnetic Resonance Research Center and n=42 participants were scanned at the Yale University Brain Imaging Center (**Table 3**). Scanning sequences were based on those used in the Adolescent Brain Cognitive Development Study^55^. Scan sequence details are reported in the **Supplement**.

#### FreeSurfer Preprocessing

T1-weighted images were submitted to FreeSurfer 6.0.0 and processed using the *recon_all* pipeline^56^. Resulting estimated total intracranial volume values for each participant were regressed from white matter tract data in subsequent modeling to address variability in white matter tract integrity related to overall head size. Estimated total intracranial volume is presented in **Table S2.**

#### QSIPrep Preprocessing

Preprocessing was performed using QSIPrep 0.14.3^57^, which utilizes Nipype 1.6.1^58^. Preprocessing details can be found in the **Supplement**. Briefly, anatomical data were corrected for intensity nonuniformity, skull-stripped, and spatially normalized using ANTS 2.3.1^59^. Brain tissue segmentation was performed using FSL’s FAST^60^. Diffusion data were denoised, corrected for ringing and B1 field inhomogeneity, and intensity-matched across B0 images using tools from MRtrix^61,62^. Next, FSL’s eddy tool was used for head motion and eddy current correction as well as outlier replacement^63^. Finally, data were resampled to 1mm isotropic voxels in ACPC space. Descriptive statistics for motion metrics produced by QSIPrep are presented in **Table S2.**

#### DSI Studio Reconstruction

Diffusion orientation distribution functions (ODFs) were reconstructed using generalized q-sampling imaging^32^ with a ratio of mean diffusion distance of 1.25. 5 million streamlines were created with a maximum length of 250mm, minimum length of 30mm, random seeding, a step size of 1mm and an automatically calculated QA threshold. Scalar images representing QA, GFA, and the isotropic ODF component were produced^57^.

#### MrTrix3 Reconstruction

Multi-tissue fiber response functions were estimated using the dhollander algorithm. Fiber orientation distributions (FODs) were estimated via constrained spherical deconvolution^61^ using an unsupervised multi-tissue method^64^. Reconstruction was done using MRtrix3^61^. FODs were intensity-normalized using mtnormalize^65^.

#### Tractography and Measures of Interest

White matter tract bundles were computed for 48 tracts using TractSeg^66^. TractSeg is a novel automated toolbox that leverages advances in convolutional neural network science to learn white matter tract mappings, and has shown high accuracy compared to more traditional methods^66^. Tracts were identified and segmented using reconstructed and normalized white matter ODF images produced by MrTrix3 reconstruction. Tract segmentations were then applied to diffusion metrics of interest (generalized fractional anisotropy images, quantitative anisotropy images, and radial diffusivity images produced by DSI Studio reconstruction) to obtain measures of white matter integrity per tract using the Tractometry implementation in TractSeg^67^. Metric-specific mean integrity values were computed across participants for each tract. For details on quality assurance, neuroimaging exclusion criteria, and outlier removal, please see the **Supplement**.

### Statistical Analysis

#### Data preparation

Analyses were conducted in python (version 3.7.12)^68^. Tables were created in R (version 4.4.2)^69^ using the package *arsenal* and python (version 3.7.12). Prior to submitting white matter and adversity exposure data to sparse canonical correlation analysis (sCCA), we regressed covariates from both sets of data. For tractwise diffusivity metrics, we regressed age at session, estimated total intracranial volume, mean framewise displacement during the DWI scan, a binary indicator of scan site, and mean integrity of the metric (fractional anisotropy, quantitative anisotropy, or radial diffusivity) using an Ordinary Least Squares model (OLS; implemented in *scipy,* version 1.7.3)^70^ separately for each tract. Tractwise descriptive statistics are presented each metric in **Table S3**. If the tract was affected by a segmentation abnormality, a categorical variable representing tract coverage differences was also included in the covariate regression model (see the **Supplement** for details). We opted to include these covariates due to the well-documented effects of age^71^ and head motion^72^ on white matter. We opted not to include sex assigned at birth as a covariate, instead accounting for estimated total intracranial volume, as a recent meta-analysis indicates that sex-associated differences in the brain may be largely explained by head size^73^. Finally, for each metric of interest (GFA, QA, and RD) we computed mean value across all tracts in the brain for each participant, and included that mean value as a covariate in a metric-specific manner in order to better account for brain-wide individual differences in integrity and to isolate tract-specific variability. For adversity exposure endorsement data, we regressed age at session and the total number of adversity exposures in adulthood (between the ages of 18-30; or the participant’s current age if they were not yet 30) per participant using a zero-inflated Poisson regression (implemented in *statsmodels,* version 0.13.5)^74^. By regressing adversity exposures in adulthood, we aimed to reduce variability related to cumulative additional exposures, thereby isolating variance specific to ages 0-18. Importantly, we intentionally regressed a separate sets of covariates from the diffusion and adversity exposure datasets, as recent work demonstrates that regression of the same set of covariates from both datasets may introduce dependencies among the datasets and inflate the likelihood of type I error^75^. For details on model and hyperparameter selection, please see the **Supplement**.

#### Model fitting

Three separate sCCA models were fitted (implemented the package *cca-zoo,* version 1.16.2)^76^. Specifically, we fitted models examining, respectively, links between the developmental timing of adversity exposure and GFA, QA, and RD. We reasoned that given that these metrics measure overlapping properties of underlying white matter but may be differentially sensitive to certain cellular properties that vary as a function of development (e.g., myelination)^32,33^, identifying measure-specific links with adversity data would yield nuanced results regarding how timing of adversity is associated with white matter tract properties. To identify the most robust patterns of association, we fitted both an unshuffled model and a shuffled (null) model for each of 10,000 iterations of resampling. For each resampling iteration, we fitted a sCCA model with 19 modes (corresponding to the smallest dimension of variables across the two datasets) and computed covariance explained and correlation strengths for each mode. Then, to evaluate model significance, we fully shuffled the adversity data (thereby unlinking the adversity and white matter tract data) and computed covariance explained and correlation strengths from the resulting null model. To evaluate the significance of the unshuffled models relative to the shuffled (null) models, we used permutation testing^77^ (i.e., counting the number of correlation strengths from the null distribution that surpassed the median unshuffled correlation strength for a given mode and dividing that number by the number of iterations (10,000). We opted to use the median (rather than the mean) as a representative metric as the correlation values did not follow a Gaussian distribution. We then corrected for multiple comparisons using FDR correction^36^. That is, we selected modes that explained more than 10% of the overall covariance and applied FDR correction to the p-values for each mode resulting from permutation testing.

#### Associations with Symptoms

To test whether our model identified combinations of adversity and white matter tract measures that were relevant to mental health, for each model fit across 10,000 iterations of resampling, we subsequently transformed the input data using the fitted model, resulting in participant-level values for adversity and diffusion variates across all fitted modes. Following the completion of all iterations of model fitting, we computed mean participant-level values for the adversity and white matter variates of the first mode. In line with our previous decision to focus solely on the first mode, here we similarly examined only the transformed variate values of adversity and white matter tract integrity of the first mode in the GFA, QA, and RD models. We tested whether measures of total psychopathology symptoms (total symptoms T-scores derived from the Adult Self Report scale and trauma-related symptoms derived from the TSC-40) were associated with sCCA model variates by fitting one OLS model (implemented in *statsmodels,* version 0.13.5)^74^ per symptom measure (i.e., scales from the ASR and TSC-40) for the adversity variates of the three models (GFA, QA, and RD) and a separate OLS model for the diffusion variates of three models. We used FDR correction to account for multiple comparisons across the two symptom measures. We opted to test white matter and adversity variates in separate models since by design sCCA identifies modes representing adversity and white matter variates that are highly correlated with each other and including them within the same model could destabilize effect estimations. In all models, we used the Variance Inflation Factor (VIF) to confirm that the sCCA-derived variates were not problematically collinear (all VIFs < 2), and the Jarque-Bera test to confirm that our measure of psychopathology symptoms followed a normal distribution. We found that while total symptoms T-scores approximated a normal distribution (JB = 4.568, *p* = .102), trauma-related symptoms did not (JB = 26.715, *p* < .001). In order to ensure that the measure met linear model assumptions, we applied a square root transformation to the trauma-related symptoms measure, which resulted in an approximately normal distribution (JB = 1.702, *p* = .427). The transformed trauma-related symptoms measure was used in all subsequent modeling.

#### Sensitivity Analyses

Next, we conducted sensitivity analyses to probe whether associations between model variates and mental health symptoms were related to proxies of socioeconomic status (total family income and years of education). Specifically, we refitted the OLS models, including total family income and years of education as covariates. To test whether the variables that loaded most strongly onto each variate were independently associated with symptoms, we fitted a separate OLS model for each of the top 25% loading variables of each variate, respectively (i.e., adversity and tract integrity), to see whether these variables were independently associated with symptoms.

#### Data and Code availability

Code used to run QSIPrep preprocessing and reconstruction and TractSeg tractography can be found at https://github.com/Yale-CANDLab/DWI_Code. The dataset and the code used to prepare and analyze the dataset can be found at https://github.com/Yale-CANDLab/Sisk_DWI_Adversity_sCCA.

**Figure 4.**
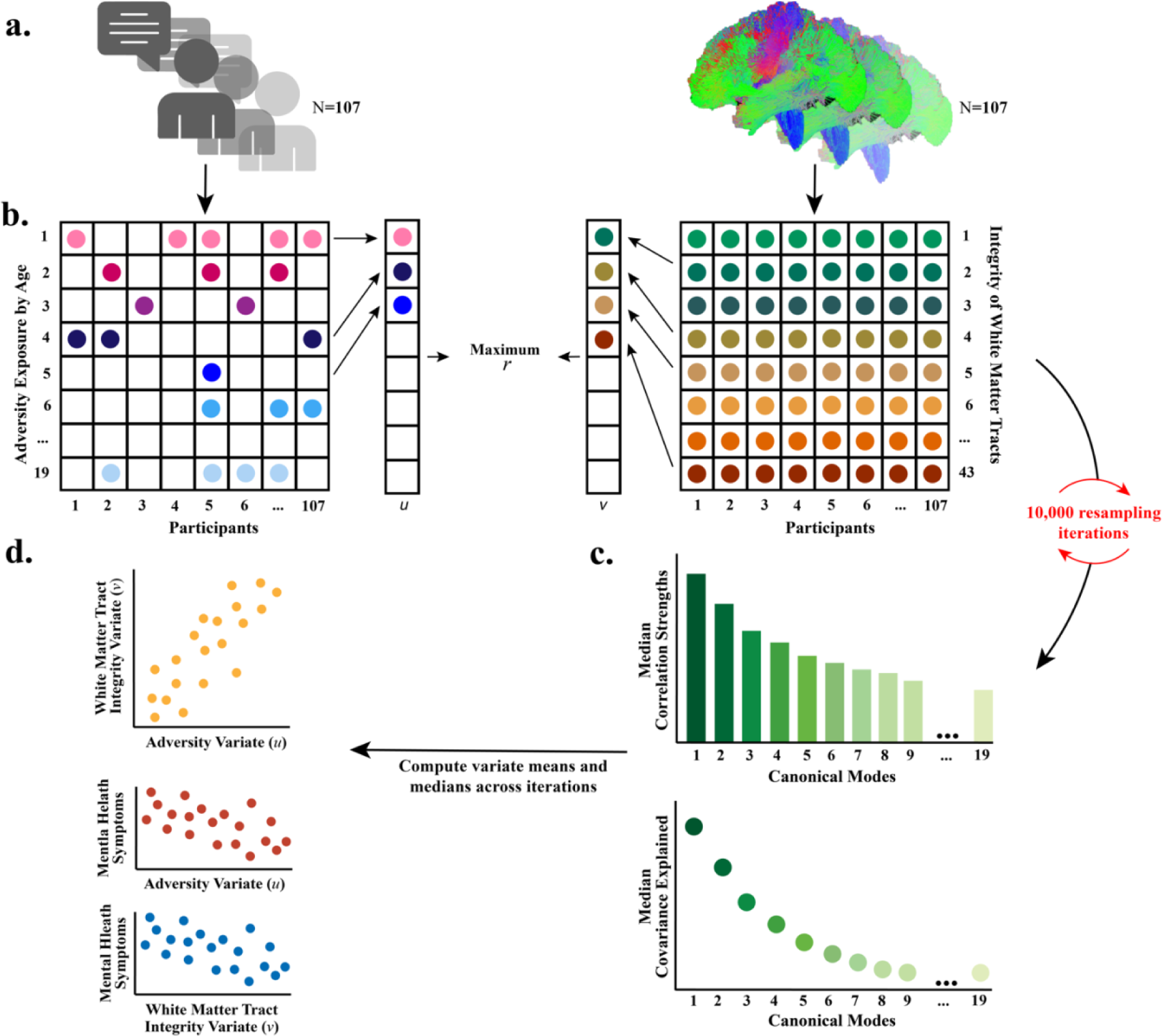
Schematic of approach. **a.** Dimensional adversity exposure data from the DISTAL (left) and white matter tract integrity measures from 43 tracts (right) were submitted to sparse canonical correlation analysis (sCCA). **b.** Nineteen sCCA modes were fitted per sCCA model, and this process was repeated 10,000 times with resampling. For each iteration, a shuffled (null) model was also fitted. **c.** For each iteration, correlation strengths for each mode and covariance explained for each mode were retained, as were variable loadings for each variate of each mode. **d.** sCCA variates maximize correlation strengths. Following sCCA modeling, associations between transformed participant-level variates and mental health symptoms (assessed via the ASR and TSC-40) were tested.

## Author Contributions

DGG, LMS, EMC, and PO contributed to study design. LMS and DGG conceptualized and designed the analyses for this project. LMS and TJK analyzed the data. EMC, SM, JCP, PO, SK, JTH, SJZ, HRH, LMS, and CC contributed to data collection. LMS, GG, AYH, AT, EMC, SM, JCP, and HRH contributed to quality assessment and data validation. LMS drafted the paper and TJK, EMC, PO, SK, SJZ, HRH, GG, AYH, AT, and DGG provided review and editing.

## Acknowledgements

This work was supported by funding from National Science Foundation (NSF) Graduate Research Fellowship Program award (NSF DGE-1752134) to LMS; Yale Child Study Center Postdoctoral T32MH18268 and Brain & Behavior Research Foundation (NARSAD) Young Investigator Award #28436 to TJK; NSF Graduate Research Fellowship Program award (NSF DGE-1752134), American Psychological Foundation Elizabeth Munsterberg Koppitz Child Psychology Graduate Fellowship, The Society for Clinical Child and Adolescent Psychology (Division 53 of the American Psychological Association) Donald Routh Dissertation Grant, a Dissertation Funding Award from the Society for Research in Child Development, a Dissertation Research Award from the American Psychological Association, an American Dissertation Fellowship from the American Association of University Women (AAUW), and a Scholar Award granted by the International Chapter of the Philanthropic educational Organization (P.E.O. Foundation) to EMC; NSF Graduate Research Fellowship Program award (NSF DGE-1752134) to PO; Janet and Sheldon (1959) Razin Fellowship (MIT) to SJZ; NSF CAREER Award (BCS-2145372), National Institutes of Health (NIH) Director’s Early Independence Award (DP5OD021370), Brain & Behavior Research Foundation (NARSAD) Young Investigator Award, Jacobs Foundation Early Career Research Fellowship, and The Society for Clinical Child and Adolescent Psychology (Division 53 of the American Psychological Association) Richard “Dick” Abidin Early Career Award and Grant to DGG.

## Notes

https://github.com/Yale-CANDLab/Sisk_DWI_Adversity_sCCA

https://github.com/Yale-CANDLab/DWI_Code

### Competing Interest Statement

The authors have declared no competing interest.

