## Supplement for "Multivariate links between the developmental timing of adversity exposure and white matter tract integrity in adulthood"

**Supplemental Methods.**

***Exclusion Criteria.*** Additional exclusion criteria included 1) history of head injury or concussion, 2) history of chronic medical illness or neurological disorder, 3) lifetime history of psychotic disorders, autism spectrum disorder, bipolar disorder, conduct disorder, non-alcohol or non-tobacco substance use disorder, primary current diagnosis of attention-deficit/hyperactivity disorder or major depressive disorder, 4) current alcohol or tobacco substance use disorder, 5) acute suicidal ideation, 6) current use of psychotropic medication, 7) colorblindness, 8) visual impairment that cannot be corrected, and 9) hearing impairment.

***Scan Sequence Parameters.***

A whole-brain high-resolution T1-weighted anatomical scan with magnetization-prepared rapid acquisition gradient echo (MPRAGE; 1070 ms TI, 2500 ms TR; 2.9 ms TE; 8° flip angle; 256 mm field of view (FoV); 176 slices in sagittal plane; 256 × 256 matrix; 2× parallel imaging; 1.0 × 1.0 × 1.0 mm resolution) was collected, as was a high angular resolution diffusion imaging (HARDI) scan, with multiple b-values, and fast integrated B0 distortion correction (reversed polarity gradient (RPG) method^1,2^ (4100 ms TR, 88 ms TE, 90° flip angle, 240 mm FoV, 81 slices in sagittal plane, 140 x 140 matrix, 1.7 x 1.7 x 1.7 mm resolution). Reverse phase-encoded fieldmaps were also collected immediately prior to the HARDI sequence.

***Preprocessing and Reconstruction.***

*Anatomical data preprocessing:* The T1-weighted (T1w) image was corrected for intensity non-uniformity (INU) using N4BiasFieldCorrection (ANTs 2.3.1)^3^, and used as T1w-reference throughout the workflow. The T1w-reference was then skull-stripped using antsBrainExtraction.sh (ANTs 2.3.1), using OASIS as target template. Spatial normalization to the ICBM 152 Nonlinear Asymmetrical template version 2009c^4^ was performed through nonlinear registration with antsRegistration (ANTs 2.3.1, RRID:SCR_004757)^5^, using brain-extracted versions of both T1w volume and template. Brain tissue segmentation of cerebrospinal fluid (CSF), white-matter (WM) and gray-matter (GM) was performed on the brain-extracted T1w using FAST (FSL 6.0.3:b862cdd5, RRID:SCR_002823)^6^.

*Diffusion data preprocessing*. Any images with a b-value less than 100 s/mm^2^ were treated as a b=0 image. MP-PCA denoising as implemented in MRtrix3’s dwidenoise^7^ was applied with a 5-voxel window. After MP-PCA, Gibbs unringing was performed using MRtrix3’s mrdegibbs^8^. Following unringing, B1 field inhomogeneity was corrected using dwibiascorrect from MRtrix3 with the N4 algorithm^3^. After B1 bias correction, the mean intensity of the DWI series was adjusted so all the mean intensity of the b=0 images matched across each separate DWI scanning sequence. FSL (version 6.0.3b862cdd5)’s eddy was used for head motion correction and Eddy current correction^9^. Eddy was configured with a q-space smoothing factor of 10, a total of 5 iterations, and 1000 voxels used to estimate hyperparameters. A linear first-level model and a linear second-level model were used to characterize Eddy current-related spatial distortion. q-space coordinates were forcefully assigned to shells. Field offset was attempted to be separated from participant movement. Shells were aligned post-eddy. Eddy’s outlier replacement was run^10^. Data were grouped by slice, only including values from slices determined to contain at least 250 intracerebral voxels. Groups deviating by more than 4 standard deviations from the prediction had their data replaced with imputed values. Data were collected with reversed phase-encode blips, resulting in pairs of images with distortions going in opposite directions. Here, b=0 reference images with reversed phase encoding directions were used along with an equal number of b=0 images extracted from the DWI scans. From these pairs the susceptibility-induced off-resonance field was estimated using a method similar to that described in^11^. The fieldmaps were ultimately incorporated into the Eddy current and head motion correction interpolation. Final interpolation was performed using the jac method. Several confounding time-series were calculated based on the preprocessed DWI: framewise displacement (FD) using the implementation in Nipype^12^. The head-motion estimates calculated in the correction step were also placed within the corresponding confounds file (**Table S2**). Slicewise cross correlation was also calculated. The DWI time-series were resampled to ACPC space, generating a preprocessed DWI run in ACPC space with upsampled 1mm isotropic voxels. Upsampling has been shown to improve registration accuracy^13^. Many internal operations of QSIPrep use Nilearn 0.8.1 (Abraham et al., 2014, RRID:SCR_001362) and Dipy^15^. For more details of the pipeline, see the section corresponding to workflows in QSIPrep’s documentation.

*Quality assurance.* Raw diffusion weighted imaging (DWI) scans were manually inspected for artifacts. After preprocessing, reports generated by QSIprep that included images from each processing step were also inspected per participant to ensure registration accuracy, brain mask accuracy, and proper alignment. All cerebellar tracts were excluded from analyses, as 27 participants (25% of sample) had cerebellar cutoff due to scan prescription, resulting in 5 of the 48 tracts generated by TractSeg being excluded across all metrics. Specifically, we excluded bilateral inferior cerebellar peduncle tracts, bilateral superior cerebellar peduncle tracts, and the medial cerebellar peduncle tract. The remaining 43 tracts were visually inspected for anatomical accuracy. Several tract segmentations (specifically the corpus callosum rostral body, the right superior longitudinal fasciculus I, the left superior longitudinal fasciculus II, and the left superior thalamic radiation) displayed differences in size and regional coverage; a categorical variable representing coverage differences across participants was created and regressed across participants as a nuisance covariate from tract integrity data. See Supplementary Information for an image and detailed description.

*Exclusion criteria and outlier removal.* 124 participants had DWI scan data and completed the DISTAL interview. Motion metrics produced by QSIPrep (mean framewise displacement, maximum framewise displacement, maximum rotation, maximum relative rotation, maximum translation, maximum relative translation, and number of bad slices after processing) were used to evaluate data quality. Participants whose values for any of these metrics were more than 3 standard deviations from the mean were excluded (n=6). See **Table S2** for motion descriptive statistics. Participants who met diagnostic exclusion criteria were also excluded from the sample (n=4). Participants with anomalies detected at the scan session (n=2), participants who were missing data for age at session (n=1), and participants with adversity exposure endorsement counts that were more than 3 standard deviations from the median for one or more years of age (n=4) were also excluded. This yielded the final sample size of 107 participants.

*Model and hyperparameter selection.* We opted to perform our analysis using sparse canonical correlation analysis (sCCA)^16,17^, a model which identifies linear combinations of variables that maximize covariance explained between two sets of high-dimensional data. The model was implemented in python (version 3.7.12) in the package *cca-zoo* (version 1.16.2)^18^. A full list of package versions can be found in the project Github repository. To balance the richness of information provided by the developmentally-informed DISTAL while simultaneously limiting overfitting through regularization, we opted to use hyperparameters enforcing moderate sparsity across both sets of data (C=0.5 for both adversity and white matter datasets) rather than identifying hyperparameters through a grid search.

**Table S1.**

|  | **Overall (N=107)** |
| --- | --- |
| **Ages 0-1 (n=6)** |  |
| Mean (SD) | 1.000 (0.000) |
| Range | 1.000 - 1.000 |
| **Ages 1-2 (n=5)** |  |
| Mean (SD) | 1.200 (0.447) |
| Range | 1.000 - 2.000 |
| **Ages 2-3 (n=9)** |  |
| Mean (SD) | 1.111 (0.333) |
| Range | 1.000 - 2.000 |
| **Ages 3-4 (n=8)** |  |
| Mean (SD) | 1.250 (0.707) |
| Range | 1.000 - 3.000 |
| **Ages 4-5 (n=13)** |  |
| Mean (SD) | 1.231 (0.599) |
| Range | 1.000 - 3.000 |
| **Ages 5-6 (n=23)** |  |
| Mean (SD) | 1.348 (0.647) |
| Range | 1.000 - 3.000 |
| **Ages 6-7 (n=29)** |  |
| Mean (SD) | 1.414 (0.682) |
| Range | 1.000 - 3.000 |
| **Ages 7-8 (n=30)** |  |
| Mean (SD) | 1.467 (0.937) |
| Range | 1.000 - 4.000 |
| **Ages 8-9 (n=39)** |  |
| Mean (SD) | 1.590 (0.938) |
| Range | 1.000 - 5.000 |
| **Ages 9-10 (n=31)** |  |
| Mean (SD) | 1.742 (1.341) |
| Range | 1.000 - 7.000 |
| **Ages 10-11 (n=44)** |  |
| Mean (SD) | 1.659 (1.098) |
| Range | 1.000 - 6.000 |
| **Ages 11-12 (n=43)** |  |
| Mean (SD) | 1.698 (1.013) |
| Range | 1.000 - 6.000 |
| **Ages 12-13 (n=55)** |  |
| Mean (SD) | 2.182 (1.529) |
| Range | 1.000 - 6.000 |
| **Ages 13-14 (55)** |  |
| Mean (SD) | 1.891 (1.100) |
| Range | 1.000 - 6.000 |
| **Ages 14-15 (n=58)** |  |
| Mean (SD) | 1.983 (1.192) |
| Range | 1.000 - 6.000 |
| **Ages 15-16 (n=57)** |  |
| Mean (SD) | 2.281 (1.601) |
| Range | 1.000 - 7.000 |
| **Ages 16-17 (n=61)** |  |
| Mean (SD) | 2.623 (1.781) |
| Range | 1.000 - 8.000 |
| **Ages 17-18 (n=72)** |  |
| Mean (SD) | 2.097 (1.313) |
| Range | 1.000 - 7.000 |
| **Ages 18-19 (n=67)** |  |
| Mean (SD) | 2.418 (1.568) |
| Range | 1.000 - 7.000 |
| **Ages 19-20 (n=64)** |  |
| Mean (SD) | 2.250 (1.321) |
| Range | 1.000 - 6.000 |
| **Ages 20-21 (n=48)** |  |
| Mean (SD) | 2.167 (1.117) |
| Range | 1.000 - 5.000 |
| **Ages 21-22 (n=44)** |  |
| Mean (SD) | 2.409 (1.452) |
| Range | 1.000 - 7.000 |
| **Ages 22-23 (n=38)** |  |
| Mean (SD) | 2.211 (1.580) |
| Range | 1.000 - 7.000 |
| **Ages 23-24 (25)** |  |
| Mean (SD) | 1.920 (1.038) |
| Range | 1.000 - 4.000 |
| **Ages 24-25 (n=23)** |  |
| Mean (SD) | 2.087 (1.411) |
| Range | 1.000 - 5.000 |
| **Ages 25-26 (n=16)** |  |
| Mean (SD) | 2.312 (1.852) |
| Range | 1.000 - 7.000 |
| **Ages 26-27 (n=12)** |  |
| Mean (SD) | 2.667 (1.435) |
| Range | 1.000 - 6.000 |
| **Ages 27-28 (n=3)** |  |
| Mean (SD) | 1.333 (0.577) |
| Range | 1.000 - 2.000 |
| **Ages 28-29 (n=4)** |  |
| Mean (SD) | 1.750 (0.957) |
| Range | 1.000 - 3.000 |
| **Ages 29-30 (n=2)** |  |
| Mean (SD) | 3.000 (2.828) |
| Range | 1.000 - 5.000 |
| **No age reported (n=13)** |  |
| Mean (SD) | 2.538 (3.178) |
| Range | 1.000 - 12.000 |

Table S1. For each year of age, the number of participants who endorsed at least one adverse event during this year is shown. The mean number of events during this year, the standard deviation, and the range of event counts in this year are also shown.

**Table S2.**

| **Measure** | **Overall (N=107)** |
| --- | --- |
| **eTIV (mm^3)** |  |
| Mean (SD) | 1513902.502 (189194.116) |
| Range | 904312.971 - 2038064.044 |
| **Mean Framewise Displacement (mm)** |  |
| Mean (SD) | 0.467 (0.158) |
| Range | 0.089 - 0.854 |
| **Maximum Framewise Displacement (mm)** |  |
| Mean (SD) | 1.182 (0.460) |
| Range | 0.137 - 2.247 |
| **Maximum Rotation (mm)** |  |
| Mean (SD) | 0.009 (0.005) |
| Range | 0.001 - 0.031 |
| **Maximum Relative Rotation (mm)** |  |
| Mean (SD) | 0.004 (0.002) |
| Range | 0.001 - 0.014 |
| **Maximum Translation (mm)** |  |
| Mean (SD) | 0.824 (0.342) |
| Range | 0.074 - 1.646 |
| **Maximum Relative Translation (mm)** |  |
| Mean (SD) | 0.876 (0.421) |
| Range | 0.049 - 1.993 |

*Table S2. Motion statistics derived from QSIPrep and estimated total intracranial volume derived from Freesurfer.*

**Table S3.**

|  | **GFA (N=107)** | **QA (N=107)** | **RD (N=107)** |
| --- | --- | --- | --- |
| **AF_left** |  |  |  |
| Mean (SD) | 0.097 (0.003) | 0.128 (0.026) | 0.402 (0.016) |
| Range | 0.088 - 0.106 | 0.032 - 0.195 | 0.368 - 0.473 |
| **AF_right** |  |  |  |
| Mean (SD) | 0.093 (0.003) | 0.123 (0.025) | 0.404 (0.016) |
| Range | 0.085 - 0.104 | 0.030 - 0.177 | 0.367 - 0.452 |
| **ATR_left** |  |  |  |
| Mean (SD) | 0.086 (0.003) | 0.104 (0.022) | 0.443 (0.013) |
| Range | 0.078 - 0.093 | 0.023 - 0.153 | 0.405 - 0.485 |
| **ATR_right** |  |  |  |
| Mean (SD) | 0.086 (0.003) | 0.102 (0.021) | 0.441 (0.013) |
| Range | 0.080 - 0.095 | 0.024 - 0.148 | 0.403 - 0.481 |
| **CC_1** |  |  |  |
| Mean (SD) | 0.114 (0.005) | 0.124 (0.028) | 0.427 (0.020) |
| Range | 0.101 - 0.131 | 0.028 - 0.192 | 0.365 - 0.476 |
| **CC_2** |  |  |  |
| Mean (SD) | 0.111 (0.004) | 0.135 (0.030) | 0.422 (0.020) |
| Range | 0.103 - 0.123 | 0.031 - 0.195 | 0.380 - 0.471 |
| **CC_3** |  |  |  |
| Mean (SD) | 0.112 (0.005) | 0.131 (0.028) | 0.408 (0.027) |
| Range | 0.100 - 0.124 | 0.033 - 0.197 | 0.355 - 0.474 |
| **CC_4** |  |  |  |
| Mean (SD) | 0.115 (0.004) | 0.145 (0.031) | 0.408 (0.020) |
| Range | 0.106 - 0.122 | 0.039 - 0.225 | 0.362 - 0.482 |
| **CC_5** |  |  |  |
| Mean (SD) | 0.118 (0.004) | 0.144 (0.031) | 0.436 (0.025) |
| Range | 0.109 - 0.128 | 0.038 - 0.217 | 0.381 - 0.502 |
| **CC_6** |  |  |  |
| Mean (SD) | 0.122 (0.004) | 0.154 (0.031) | 0.400 (0.019) |
| Range | 0.114 - 0.133 | 0.039 - 0.232 | 0.361 - 0.476 |
| **CC_7** |  |  |  |
| Mean (SD) | 0.132 (0.007) | 0.166 (0.034) | 0.424 (0.032) |
| Range | 0.114 - 0.145 | 0.044 - 0.258 | 0.359 - 0.516 |
| **CG_left** |  |  |  |
| Mean (SD) | 0.099 (0.005) | 0.118 (0.025) | 0.427 (0.018) |
| Range | 0.088 - 0.115 | 0.028 - 0.183 | 0.373 - 0.473 |
| **CG_right** |  |  |  |
| Mean (SD) | 0.099 (0.004) | 0.115 (0.024) | 0.418 (0.017) |
| Range | 0.090 - 0.112 | 0.029 - 0.182 | 0.369 - 0.464 |
| **CST_left** |  |  |  |
| Mean (SD) | 0.111 (0.003) | 0.146 (0.030) | 0.370 (0.012) |
| Range | 0.103 - 0.119 | 0.038 - 0.217 | 0.342 - 0.407 |
| **CST_right** |  |  |  |
| Mean (SD) | 0.111 (0.004) | 0.146 (0.031) | 0.371 (0.013) |
| Range | 0.103 - 0.120 | 0.038 - 0.223 | 0.337 - 0.403 |
| **FPT_left** |  |  |  |
| Mean (SD) | 0.103 (0.003) | 0.128 (0.027) | 0.389 (0.012) |
| Range | 0.095 - 0.110 | 0.033 - 0.191 | 0.357 - 0.418 |
| **FPT_right** |  |  |  |
| Mean (SD) | 0.104 (0.004) | 0.128 (0.027) | 0.388 (0.012) |
| Range | 0.096 - 0.114 | 0.033 - 0.190 | 0.353 - 0.413 |
| **IFO_left** |  |  |  |
| Mean (SD) | 0.107 (0.004) | 0.130 (0.026) | 0.442 (0.017) |
| Range | 0.096 - 0.116 | 0.032 - 0.184 | 0.411 - 0.487 |
| **IFO_right** |  |  |  |
| Mean (SD) | 0.107 (0.003) | 0.126 (0.025) | 0.446 (0.016) |
| Range | 0.098 - 0.115 | 0.031 - 0.184 | 0.407 - 0.503 |
| **ILF_left** |  |  |  |
| Mean (SD) | 0.105 (0.004) | 0.126 (0.026) | 0.444 (0.018) |
| Range | 0.094 - 0.114 | 0.030 - 0.178 | 0.400 - 0.490 |
| **ILF_right** |  |  |  |
| Mean (SD) | 0.104 (0.004) | 0.124 (0.025) | 0.441 (0.019) |
| Range | 0.096 - 0.113 | 0.031 - 0.178 | 0.400 - 0.491 |
| **OR_left** |  |  |  |
| Mean (SD) | 0.107 (0.004) | 0.136 (0.028) | 0.430 (0.017) |
| Range | 0.097 - 0.116 | 0.034 - 0.210 | 0.393 - 0.469 |
| **OR_right** |  |  |  |
| Mean (SD) | 0.107 (0.004) | 0.139 (0.029) | 0.427 (0.018) |
| Range | 0.100 - 0.117 | 0.035 - 0.218 | 0.383 - 0.501 |
| **POPT_left** |  |  |  |
| Mean (SD) | 0.108 (0.003) | 0.139 (0.028) | 0.400 (0.015) |
| Range | 0.101 - 0.117 | 0.037 - 0.208 | 0.365 - 0.447 |
| **POPT_right** |  |  |  |
| Mean (SD) | 0.110 (0.003) | 0.140 (0.029) | 0.404 (0.017) |
| Range | 0.102 - 0.117 | 0.038 - 0.207 | 0.366 - 0.454 |
| **SLF_I_left** |  |  |  |
| Mean (SD) | 0.099 (0.004) | 0.131 (0.028) | 0.405 (0.014) |
| Range | 0.090 - 0.108 | 0.034 - 0.195 | 0.369 - 0.446 |
| **SLF_I_right** |  |  |  |
| Mean (SD) | 0.099 (0.004) | 0.130 (0.028) | 0.401 (0.015) |
| Range | 0.091 - 0.108 | 0.035 - 0.192 | 0.367 - 0.458 |
| **SLF_II_left** |  |  |  |
| Mean (SD) | 0.089 (0.004) | 0.120 (0.025) | 0.407 (0.015) |
| Range | 0.079 - 0.096 | 0.030 - 0.182 | 0.376 - 0.465 |
| **SLF_II_right** |  |  |  |
| Mean (SD) | 0.091 (0.004) | 0.121 (0.025) | 0.401 (0.016) |
| Range | 0.080 - 0.101 | 0.030 - 0.181 | 0.366 - 0.469 |
| **SLF_III_left** |  |  |  |
| Mean (SD) | 0.100 (0.004) | 0.125 (0.026) | 0.401 (0.016) |
| Range | 0.090 - 0.109 | 0.030 - 0.196 | 0.363 - 0.457 |
| **SLF_III_right** |  |  |  |
| Mean (SD) | 0.098 (0.003) | 0.121 (0.025) | 0.405 (0.016) |
| Range | 0.090 - 0.107 | 0.029 - 0.180 | 0.363 - 0.446 |
| **STR_left** |  |  |  |
| Mean (SD) | 0.094 (0.004) | 0.124 (0.026) | 0.400 (0.012) |
| Range | 0.085 - 0.103 | 0.033 - 0.183 | 0.367 - 0.435 |
| **STR_right** |  |  |  |
| Mean (SD) | 0.095 (0.004) | 0.124 (0.026) | 0.398 (0.011) |
| Range | 0.087 - 0.106 | 0.034 - 0.180 | 0.368 - 0.427 |
| **UF_left** |  |  |  |
| Mean (SD) | 0.090 (0.004) | 0.104 (0.022) | 0.478 (0.019) |
| Range | 0.077 - 0.102 | 0.024 - 0.151 | 0.420 - 0.528 |
| **UF_right** |  |  |  |
| Mean (SD) | 0.089 (0.004) | 0.100 (0.021) | 0.481 (0.018) |
| Range | 0.079 - 0.099 | 0.025 - 0.144 | 0.433 - 0.519 |
| **T_PREM_left** |  |  |  |
| Mean (SD) | 0.090 (0.003) | 0.109 (0.022) | 0.409 (0.012) |
| Range | 0.081 - 0.100 | 0.028 - 0.166 | 0.375 - 0.440 |
| **T_PREM_right** |  |  |  |
| Mean (SD) | 0.088 (0.003) | 0.106 (0.022) | 0.410 (0.012) |
| Range | 0.080 - 0.097 | 0.027 - 0.155 | 0.365 - 0.440 |
| **T_PAR_left** |  |  |  |
| Mean (SD) | 0.098 (0.003) | 0.126 (0.025) | 0.420 (0.015) |
| Range | 0.091 - 0.105 | 0.032 - 0.185 | 0.386 - 0.483 |
| **T_PAR_right** |  |  |  |
| Mean (SD) | 0.098 (0.003) | 0.125 (0.026) | 0.422 (0.016) |
| Range | 0.089 - 0.105 | 0.033 - 0.184 | 0.386 - 0.500 |
| **ST_FO_left** |  |  |  |
| Mean (SD) | 0.084 (0.005) | 0.094 (0.021) | 0.468 (0.017) |
| Range | 0.073 - 0.095 | 0.022 - 0.140 | 0.420 - 0.525 |
| **ST_FO_right** |  |  |  |
| Mean (SD) | 0.085 (0.004) | 0.092 (0.019) | 0.470 (0.017) |
| Range | 0.072 - 0.093 | 0.022 - 0.133 | 0.425 - 0.507 |
| **ST_PREM_left** |  |  |  |
| Mean (SD) | 0.084 (0.003) | 0.104 (0.021) | 0.423 (0.013) |
| Range | 0.077 - 0.092 | 0.027 - 0.156 | 0.392 - 0.460 |
| **ST_PREM_right** | |  |  |
| Mean (SD) | 0.085 (0.003) | 0.103 (0.021) | 0.415 (0.013) |
| Range | 0.078 - 0.093 | 0.026 - 0.146 | 0.383 - 0.455 |

*Table S3. Tractwise values across the three metrics included in the present study. GFA = generalized fractional anisotropy, QA = quantitative anisotropy, RD = radial diffusivity. AF = arcuate fasciculus, ATR = anterior thalamic radiation, CC 1 = corpus callosum rostrum, CC 2 = corpus callosum genu, CC 3 = corpus callosum rostral body (premotor), CC 4 = corpus callosum anterior midbody (primary motor), CC 6 = corpus callosum isthmus, CC 7 = corpus callosum splenium, CG = cingulum cingulate, CST = corticospinal tract, FPT = fronto-pontine tract, IFO = inferior fronto-occipital fasciculus, OR=optic radiation, POPT = parietal occipital pontine tract, SLF I = superior longitudinal fasciculus I, SLF II = superior longitudinal fasciculus II, SLF III = superior longitudinal fasciculus III, ST FO = striato-fronto-orbital tract, STR = superior thalamic radiation, T PAR = thalamo-parietal tract, T PREM = thalamo-premotor tract, UF = uncinate fasciculus.*
